# Fast and simple analysis of MiSeq amplicon sequencing data with MetaAmp

**DOI:** 10.1101/131631

**Authors:** Xiaoli Dong, Manuel Kleiner, Christine E. Sharp, Erin Thorson, Carmen Li, Dan Liu, Marc Strous

## Abstract

Microbial community profiling by barcoded 16S rRNA gene amplicon sequencing currently has many applications in microbial ecology. The low costs of the parallel sequencing of multiplexed samples, combined with the relative ease of data processing and interpretation (compared to shotgun metagenomes) have made this an entry-level approach. Here we present the MetaAmp pipeline for processing of SSU rRNA gene and other non-coding or protein-coding amplicon sequencing data by investigators that are inexperienced with bioinformatics procedures. It accepts single-end or paired-end sequences in fasta or fastq format from various sequencing platforms. It includes read quality control, and merging of forward and reverse reads of paired-end reads. It makes use of UPARSE, Mothur, and the SILVA database for clustering, removal of chimeric reads, taxonomic classification and generation of diversity metrics. The pipeline has been validated with a mock community of known composition. MetaAmp provides a convenient web interface as well as command line interface. It is freely available at: http://ebg.ucalgary.ca/metaamp. Since its launch two years ago, MetaAmp has been used >2,800 times, by many users worldwide.

## Introduction

Microbial communities are made up of many populations and it has long been known that a meaningful assessment of microbial community structure is almost impossible with classical cloning and Sanger sequencing approaches (Curtis et al., 2002). Therefore, high throughput DNA sequencing of a ∼400 base pair (bp) region of the prokaryotic 16S rRNA gene has become the method of choice for characterization of microbial communities (Sogin et al., 2006). This approach consists of (1) extraction of DNA from a set of samples, (2) amplification of the ∼400 bp target region using the polymerase chain reaction (PCR) with a primer pair that targets conserved sequence elements on both sides of the amplified region, (3) barcoding of the amplicons of each sample with a short “barcode” sequence unique to each sample, and (4) high throughput, “multiplexed” sequencing of the combined amplicons from all samples in a single sequencing run. After these experimental procedures, data processing ultimately results in a list of abundances of taxa present in each sample and diversity metrics (e.g. alpha- and beta-diversity) and other statistical comparisons of samples (e.g. nonlinear multidimensional scaling).

Originally, Roche 454 pyrosequencing was the sequencing method of choice, but currently the Illumina MiSeq platform is used most frequently. This platform typically yields 25 million 2 x 300 bp paired-end reads per run, which enables parallel sequencing of up to ∼400 samples at ∼50,000 paired end reads per sample.

The consumable costs of this approach can be less than $50 per sample and an experienced scientist can process hundreds of samples per week, depending on the degree of automation available. This has enabled meaningful microbial ecology investigations of, for example, human microbiota (Yatsunenko *et al.*, 2012), oceans (Fuhrman, 2009), soils (Auffret et al., 2016), and wastewater treatment systems (Vanwolterghem et al., 2014).

The approach has more recently also been applied to metabolic genes (Herbold et al., 2015) and to eukaryotic microbes, targeting the 18S rRNA gene (Mahé et al., 2014). However, the specificity of the primer sets used for these targets remains largely untested. Most likely such surveys yield a much lower coverage of the taxonomic diversity, leading to incomplete community assessments. For prokaryotic 16S rRNA genes, the primers are also known to be imperfect, but at least this imperfection has been assessed based on the large amount of available 16S rRNA gene sequences in the databases (Klindworth et al., 2013). Imperfect primers appear to be the major source of bias of the amplicon sequencing approach (Schirmer et al., 2015).

Typical computational steps needed for data analysis consist of filtering out those sequencing reads that have a poor quality, trimming off sequencing adapters and barcodes, merging of each set of paired-end reads into a single sequence (based on overlap), and assignment of sequences to samples, using the barcodes. Next, near-identical sequences are clustered using an identity cut-off. Often 97% is used as the identity cut-off, which is the more-or-less-arbitrary, but generally accepted value of within-species sequence diversity of the 16S rRNA gene. Each cluster constitutes an “operational taxonomic unit” (OTU), which is assigned to a taxon using a database and a classification algorithm. Several databases, such as SILVA (Quast et al., 2013), RDP (Cole et al., 2014) and Greengenes (McDonald et al., 2012) for ribosomal genes, and classification algorithms, such as NCBI Blast (Camacho et al., 2009) and the Ribosomal Database Project classifier (Cole et al., 2014), have found wide use. Finally, indexes of alpha-diversity are computed for each sample, beta-diversity is calculated across samples, and many other approaches are available to compare samples, analyse the distribution of OTUs as a function of specific environmental parameters, analyse co-variation between OTUs and other hypothesis testing relevant to the study (Zhou, 2015).

Many software packages are available that perform these computations. Mothur (Schloss et al., 2009) and QIIME (Caporaso et al., 2010) are the most well-known examples. However, because these applications cater to a diverse group of users with many different specific needs, these applications present a large number of tools and options. Their proper use requires significant expertise and training.

For this reason, we developed MetaAmp, for simple, fast, command-line and web-based, push-of-a-button processing of amplicon sequencing data for casual users. MetaAmp performs all steps outlined above in an automated pipeline that uses established tools with sensible default parameters. It formats the basic results in graphs and tables for rapid interpretation. It also generates files compatible with Mothur for further analysis by advanced users. Using a series of artificial, “mock” microbial communities, we validated the approach and showed that it delivers an accurate assessment of microbial community structure. MetaAmp has been available for two years now and has found wide use worldwide. Therefore, we believe that it is a useful tool for the analysis of amplicon sequencing results.

## Materials and Methods

### MetaAmp pipeline implementation

MetaAmp is an integrated and fully automated pipeline for amplicon data analysis. MetaAmp offers both a command line and a web interface. It is written using Perl and the web interface is implemented using a standard CGI framework, HTML, and Javascript. Amplicon data analysis in MetaAmp involves several key stages (Figure 1).

**Figure 1.**
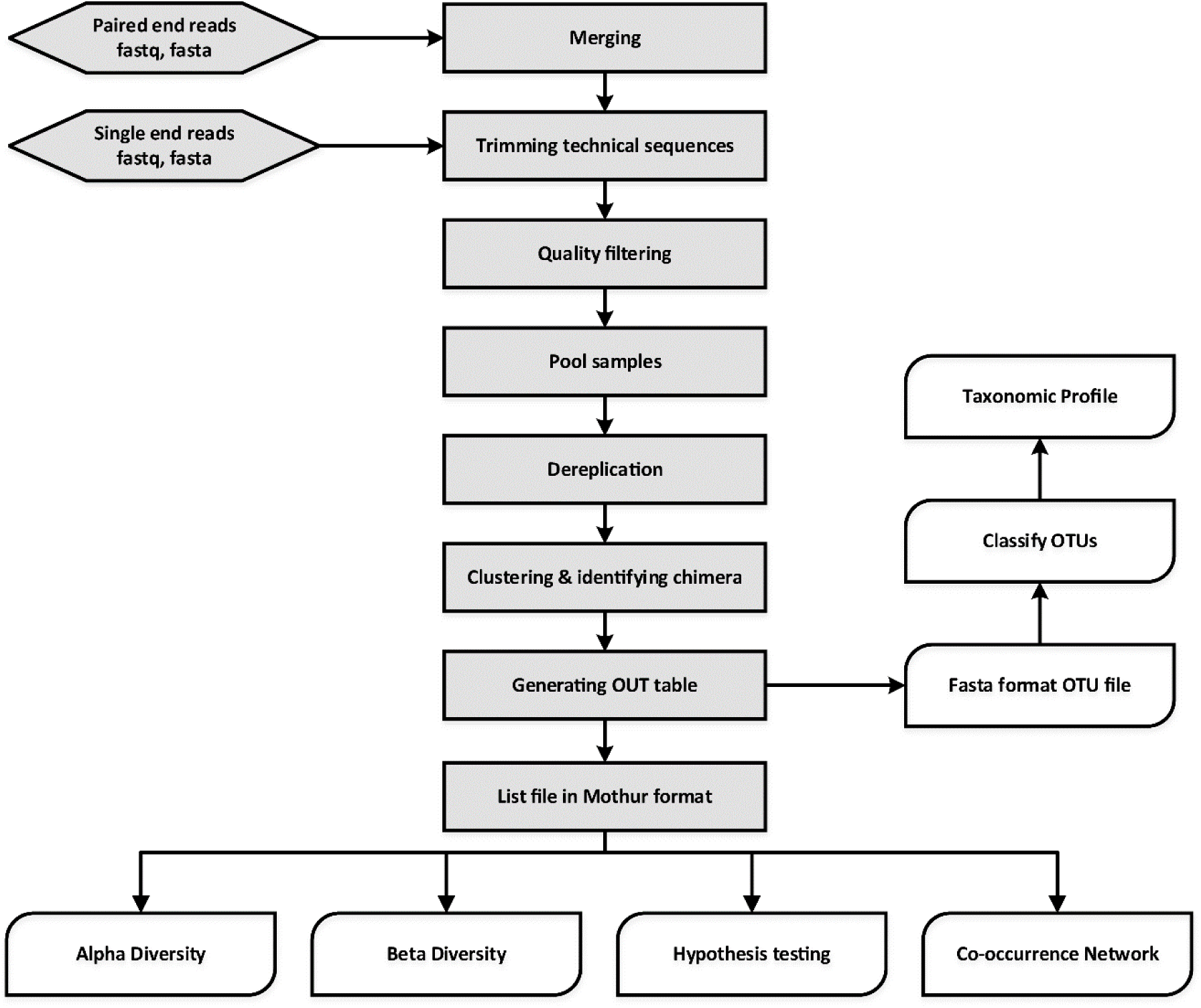
Overview of MetaAmp workflow.

The input to MetaAmp is a set of FASTA format sequence files (with matching quality files) or FASTQ format sequence files. If the input is paired-end sequence files, the paired-end raw reads from each sample will go through the assembly (merging) stage first. During assembly, a read pair is converted into a longer single read containing one sequence and one set of quality scores. A pair is merged by aligning the forward read sequence to the reverse-complement of the reverse read. In the overlap of the two reads, a single base and quality score is derived from the aligned pair of bases and quality scores for each position. MetaAmp assembles paired-end reads using “usearch - fastq_mergepairs” (Edgar, 2013). The read pairs which cannot be aligned or whose overlap regions are shorter than the user-defined length are discarded. Additionally, read pairs that have a number of mismatches in the overlap region greater than the user-defined threshold are discarded.

In a second stage, the single-end input reads or assembled reads are checked for the forward and reverse primers at the start and end of each read. The primers from both ends are trimmed with the Mothur software package (Schloss et al., 2009). Those reads which do not contain both forward and reverse primers or have a number of mismatches in the primer region greater than the user-defined threshold are discarded. The primer-trimmed reads are subjected to quality filtering to remove low-quality reads and minimize the influence of sequencing errors. The quality filtering is done using “usearch - fastq_filter”. The remaining reads are truncated to a user-defined length and reads which have a higher number of total expected errors for the truncated read length than the user-defined number are discarded. The reads with lengths shorter than the truncation length are also excluded from further analysis. After the unique sample ids are inserted into the header of the remaining high-quality reads, the reads from different files are pooled together.

In the third stage, the UPARSE software is used to dereplicate, discard singleton reads, identify chimeras and cluster the pooled high-quality reads into operational taxonomic units (OTUs). The OTU taxonomic assignments are generated using the “classify.seqs” command implemented in Mothur with the SILVA training dataset as the template (http://www.mothur.org/wiki/Taxonomy_outline). The UPARSE-generated OTU table is converted into the OTU list file format which can be used by Mothur. Since diversity and similarity measures can be highly sensitive to different sampling depths, MetaAmp rarefies each sample using the “sub.sample” command from Mothur so that rarefied samples all have the same number of reads to ensure an equal playing field for sample comparisons.

Finally, the original OTU list file and the rarefied list file are both used as input to Mothur to generate rarefaction data, rank abundance data, alpha-diversity indexes (sobs, chao, ace, jackknife, Shannon, npshannon, simpson), and beta-diversity. The community dissimilarities among different samples in terms of membership and structure are calculated using the following measures: Jclass, Jest, ThetaYC, and Bray-Curtis index. For each dissimilarity measure, the UPGMA (Unweighted Pair Group Method with Arithmetic Mean) algorithm is used to generate a Newick-format tree, which describes the hierarchical relationship among *s*amples. In addition, two ordination methods, non-metric multidimensional scaling (NMDS) and principal coordinate analysis (PCoA), are also employed to simplify and visualize the differences between microbial communities in samples. Moreover, hypothesis testing tools offered via Mothur such as parsimony, weighted unifrac and unweighted UniFrac, AMOVA, HOMOVA are used to determine the statistical significance of the spatial separation or clustering observed.

Co-occurrence relationships are ecologically important patterns that reflect niche processes that drive coexistence and diversity maintenance within biological communities (Tilman, 1982; HilleRisLambers et al., 2012, Berry et al., 2014). Network analyses-based approaches have recently been used to investigate co-occurrence patterns between microorganisms in complex environments ranging from the human gut to oceans and soils (Bin et al., 2016). MetaAmp also constructs meta-community co-occurrence networks based on Spearman or Pearson correlation coefficients and P-values. After filtering out infrequent OTUs (minimum abundance < 0.1% in the samples and only showing up in one sample), MetaAmp uses absolute abundance of the remaining OTUs to calculate Spearman and Pearson correlation coefficients between OTUs and only keeps the strong (correlation coefficient >= 0.6 and statistically significant (P-value <= 0.01 or P-value <= 0.05)) correlation pairs (Barberán et al., 2012). These correlation pairs are then used to construct a co-occurrence network in gexf file format, which can be explored and visualized in Gephi (Bastian et al., 2009).

When the MetaAmp analysis finishes, it sends the user an email notification with a web link to the main result page. The result page contains links to data files, extensive text summaries, interactive tables, and downloadable figures. The user can also download the identical packaged result set to their local computer to review the results later. For the most up to date instructions of how to use MetaAmp, please read its online help page at: http://ebg.ucalgary.ca/metaamp/html/help.html

## Testing of MetaAmp pipeline

### Construction of mock communities

Three types of mock communities were constructed as described previously (Kleiner et al. 2017). Briefly, four biological replicates of each mock community type were made by mixing between 28 and 32 species and strains of Archaea, Bacteria, Eukaryotes and Bacteriophages. Each mock community contained between 17 to 21 bacterial species that differed in their 16S rRNA gene sequence (Table 1, Figure 2). For some bacterial species, multiple strains were added that are indistinguishable on the 16S rRNA gene sequence level. These included two strains of *Rhizobium leguminosarum*, two strains of *Staphylococcus aureus*, and three strains of *Salmonella enterica* serotype typhimurium. The uneven mock community (UEC) was designed to cover a large range of species abundances on the level of cell numbers to test for the dynamic range and detection limit of the amplicon sequencing method. The equal-protein community (EPC) contained the same amount of protein for all community members. The equal-cell community (ECC) contained the same number of cells for all members.

**Figure 2.**
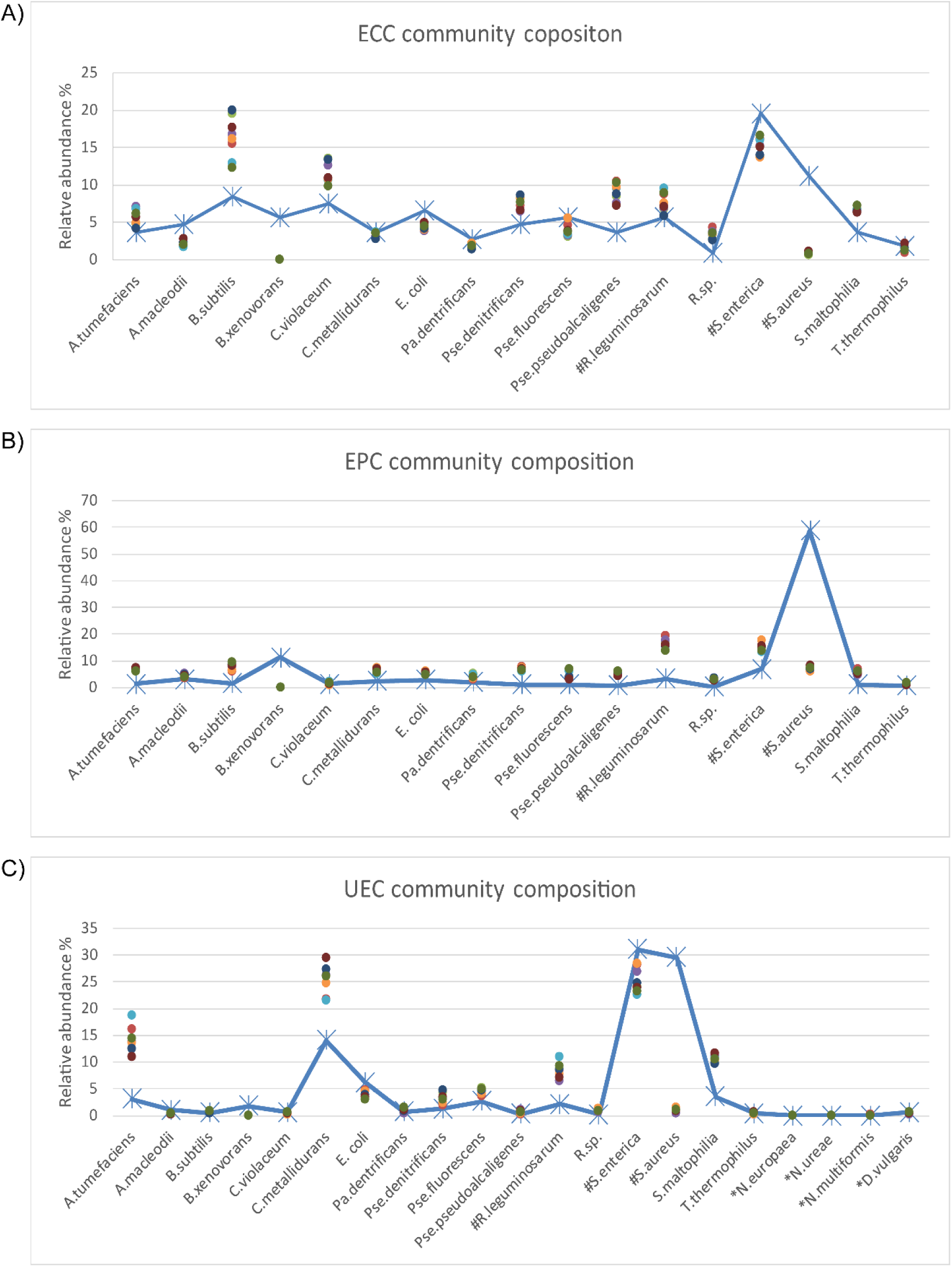
Comparison of the inferred and expected community composition. The line plots show the expected species abundance and the scatter plots show the observed abundance based on MetaAmp analysis of the MiSeq runs. Each point in the plot represents a biological or technical replicates. A) ECC is the equal cell number community. In the ECC community, the same number of cells was added to the community for each strain. B) EPC is the equal protein amount community. In the EPC community, the same amount of each strain was added to the community based on protein mass. C) UEC is the uneven community. In the UEC community, the cell count or protein amount of the different strains cover a large abundance range. The species names starting with # indicate that multiple strains were added to the mock samples and the species names starting with * indicate that those species were only added to the UEC community and their DNA input abundance was very low. The expected relative abundance of each organism was corrected by the 16s rRNA gene copy number in each organism.

**Table 1.**
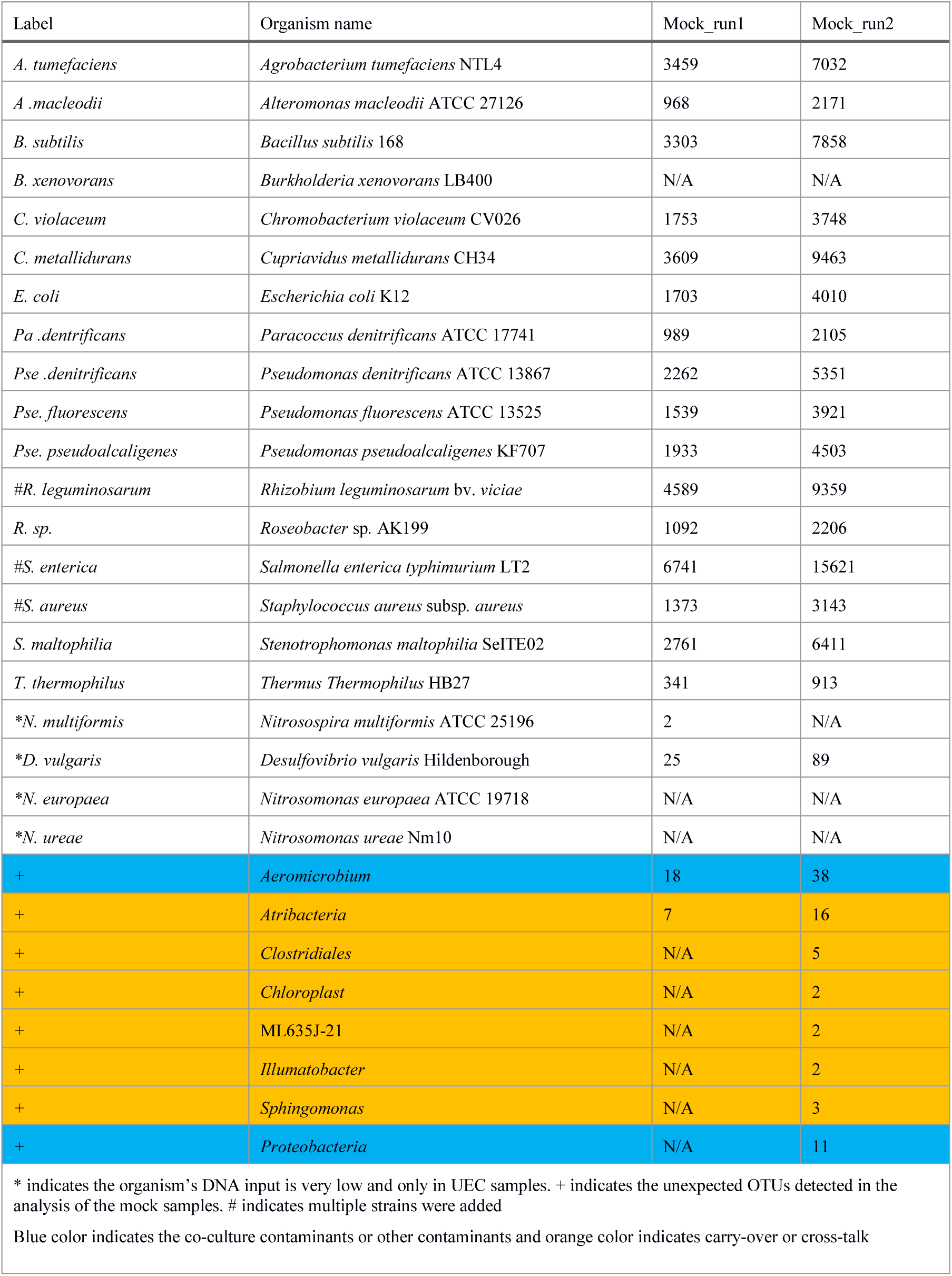
Microbial mock community input bacterial name abbreviation list and obtained community composition from the MetaAmp pipeline.

| Label | Organism name | Mock_run1 | Mock_run2 |
| --- | --- | --- | --- |
| *A. tumefaciens* | *Agrobacterium tumefaciens* NTL4 | 3459 | 7032 |
| *A. macleodii* | *Alteromonas macleodii* ATCC 27126 | 968 | 2171 |
| *B. subtilis* | *Bacillus subtilis* 168 | 3303 | 7858 |
| *B. xenovorans* | *Burkholderia xenovorans* LB400 | N/A | N/A |
| *C. violaceum* | *Chromobacterium violaceum* CV026 | 1753 | 3748 |
| *C. metallidurans* | *Cupriavidus metallidurans* CH34 | 3609 | 9463 |
| *E. coli* | *Escherichia coli* K12 | 1703 | 4010 |
| *Pa. denitrificans* | *Paracoccus denitrificans* ATCC 17741 | 989 | 2105 |
| *Pse. denitrificans* | *Pseudomonas denitrificans* ATCC 13867 | 2262 | 5351 |
| *Pse. fluorescens* | *Pseudomonas fluorescens* ATCC 13525 | 1539 | 3921 |
| *Pse. pseudoalcaligenes* | *Pseudomonas pseudoalcaligenes* KF707 | 1933 | 4503 |
| *#R. leguminosarum* | *Rhizobium leguminosarum* bv. *viciae* | 4589 | 9359 |
| *R. sp.* | *Roseobacter* sp. AK199 | 1092 | 2206 |
| *#S. enterica* | *Salmonella enterica* typhimurium LT2 | 6741 | 15621 |
| *#S. aureus* | *Staphylococcus aureus* subsp. *aureus* | 1373 | 3143 |
| *S. maltophilia* | *Stenotrophomonas maltophilia* SeITE02 | 2761 | 6411 |
| *T. thermophilus* | *Thermus Thermophilus* HB27 | 341 | 913 |
| *\*N. multiformis* | *Nitrosospira multiformis* ATCC 25196 | 2 | N/A |
| *\*D. vulgaris* | *Desulfovibrio vulgaris* Hildenborough | 25 | 89 |
| *\*N. europaea* | *Nitrosomonas europaea* ATCC 19718 | N/A | N/A |
| *\*N. ureae* | *Nitrosomonas ureae* Nm10 | N/A | N/A |
| + | *Aeromicrobium* | 18 | 38 |
| + | *Atribacteria* | 7 | 16 |
| + | *Clostridiales* | N/A | 5 |
| + | *Chloroplast* | N/A | 2 |
| + | ML635J-21 | N/A | 2 |
| + | *Illumatobacter* | N/A | 2 |
| + | *Sphingomonas* | N/A | 3 |
| + | *Proteobacteria* | N/A | 11 |
| * indicates the organism's DNA input is very low and only in UEC samples. + indicates the unexpected OTUs detected in the analysis of the mock samples. # indicates multiple strains were added<br>Blue color indicates the co-culture contaminants or other contaminants and orange color indicates carry-over or cross-talk |  |  |  |

### DNA extraction

DNA was extracted from each of the four biological replicates of each mock community type as described previously (Kleiner et al. 2017). Briefly, the FastDNA Spin Kit (MP Biomedicals, Santa Ana, CA, USA) was used according to the manufacturer’s protocol with small modifications. Samples were homogenized after addition of CLS-TC in lysing matrix tubes (MP Biomedicals FastDNA Spin Kit, tube A) for 45 seconds at 6 m/s using an OMNI Bead Ruptor 24 (Omni International, Kennesaw, GA, USA). The DNA elution step was repeated twice. DNA concentrations were measured using a NanoDrop 2000 spectrophotometer (Thermo Scientific).

### PCR amplification, barcoding and DNA sequencing

Samples were prepared for dual-index paired-end sequencing analysis as described previously (Sharp et al., 2017) using the primers S-D-Bact-0341-a-S-17 (also known as b341, 5’-

TCGTCGGCAGCGTCAGATGTGTATAAGAGACAG<u>CCTACGGGAGGCAGCAG</u>- 3’) and S-D-Bact-0785-a-A-21 (also known as Bakt_805R, 5’-

GTCTCGTGGGCTCGGAGATGTGTATAAGAGACAG<u>GACTACHVGGGTATCTAA TCC</u>-3’) with added Illumina overhang adapters on their 5’ end for the amplification of the HV regions 3-4 in the first step PCR reaction. These primers yield amplicons with a length of 427 bp and cover a large proportion of the domain Bacteria (Klindworth et al., 2013). All amplicon PCR reactions were performed in triplicate. PCR reactions were performed as described in Sharp et al. (2017). The second step PCR was used to attach dual indices and Illumina sequencing adapters to region of interest PCR products, the amplicons from the first step PCR reaction. PCR reactions and sequencing libraries were prepared as per Sharp et al., (2017) except that the magnetic DNA purification beads were sourced from Macherey-Nagel (product number 744970.500). Libraries were normalized and pooled for sequencing on the MiSeq Personal Sequencer (Illumina, San Diego, CA) using the 2 x 300 bp MiSeq Reagent Kit v3. The sequence data was deposited in EMBL-EBL under accession number ERR1938377 to ERR1938400.

### Mock sample analysis

The 12 mock samples were sequenced twice in two separate MiSeq runs. The mock sample sequence data generated from different runs were analyzed in the MetaAmp pipeline separately to create OTU taxonomic profiles, alpha-diversity and beta-diversity indexes, and hypothesis testing results with the parameters according to Table 2.

**Table 2.**
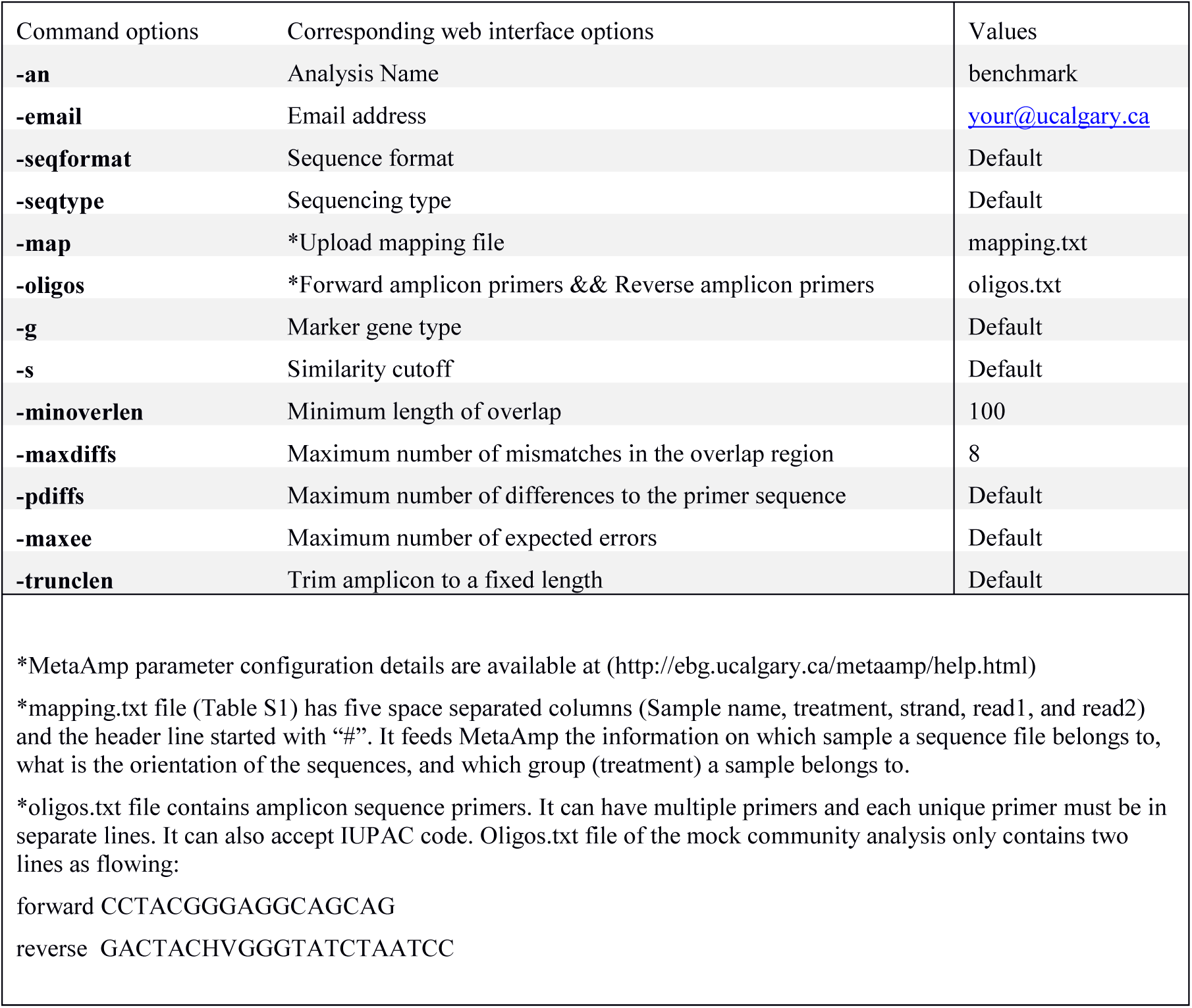
MetaAmp parameter configuration for mock community analysis

To infer the source of the non-mock community sequences in the mock samples, mock samples and all the other samples from the same and previous MiSeq runs were also analyzed in MetaAmp.

## Results

The independent analysis of mock communities from two separate MiSeq runs in MetaAmp produced two sets of analysis results referred to as Mock_run1 and Mock_run2. In both runs, MetaAmp detected 16 out of 17 bacterial species for the ECC and EPC community. MetaAmp detected 18 out of 21 bacterial species for the UEC community in Mock_run1 and 17 in Mock_run2.

MetaAmp also identified two and eight additional OTUs in Mock_run1 and Mock_run2, respectively. One of these OTUs, classified as *Aeromicrobium*, present in both runs, was a known co-cultured bacterium of the eukaryotic green algae *Chlamydomonas reinhardtii* present in the mock communities. The combined MetaAmp analysis of the mock community samples and the other samples sequenced in the same run as well as in the previous run of the MiSeq sequencer, showed that six of the unexpected OTUs from Mock_run2 and one from Mock_run1 were highly abundant in other samples sequenced in the same or the previous MiSeq run. This suggests that the detection of these OTUs resulted from carry-over or cross-talk. Sample cross-talk is a known phenomenon for multiplexed Illumina sequencing runs. At the same time, most of the abundant OTUs in the mock communities were often found in low abundance in the other samples sequenced in those runs. This presented additional evidence for cross-talk. The OTU classified as “*Sphingomonas*” is a known contaminant of commercial DNA extraction kits. The source of the remaining unexpected OTU, which yielded 11 reads in Mock_run2, could not be identified and the contamination might have occurred during library preparation or sequencing.

The representative sequences from each OTU were taxonomically classified in MetaAmp to assess how well the predicted taxonomy matched the known taxonomy of the input community. Seventeen OTUs of the mock community were correctly classified down to the genus level and NCBI blast against the mock community reference 16S rRNA gene sequences confirmed the assignment. One OTU was classified to *Rhizobium* instead of *Agrobacterium* at the genus level because *Agrobacterium* was missing from the classification reference template we downloaded from the Mothur website. However, the assignment was correct at the family level.

The MetaAmp-inferred community structures mostly agreed with the known community structures. However, several differences were observed (Figure 3). *B. xenovorans* was not detected in any of the mock communities sequenced in either MiSeq run. The inferred abundance of *S. aureus* was consistently lower than the anticipated abundance in all samples. The cause of this could be cell lysis efficiency, DNA extraction-, primer-, or PCR-bias. Three organisms (*N.multiformis*, *N.ureae*, and *N.europaea*), which were absent in Mock_run1 and/or Mock_run2 (Table 2), all had very low expected abundances (∼0.01%). Because of the compositional nature of these data, the absence or underestimation of these OTUs led to slight differences between the actual and observed abundance for all the other taxa. At the same time, all biological and technical replicates of each type of the mock community were consistent.

**Figure 3.**
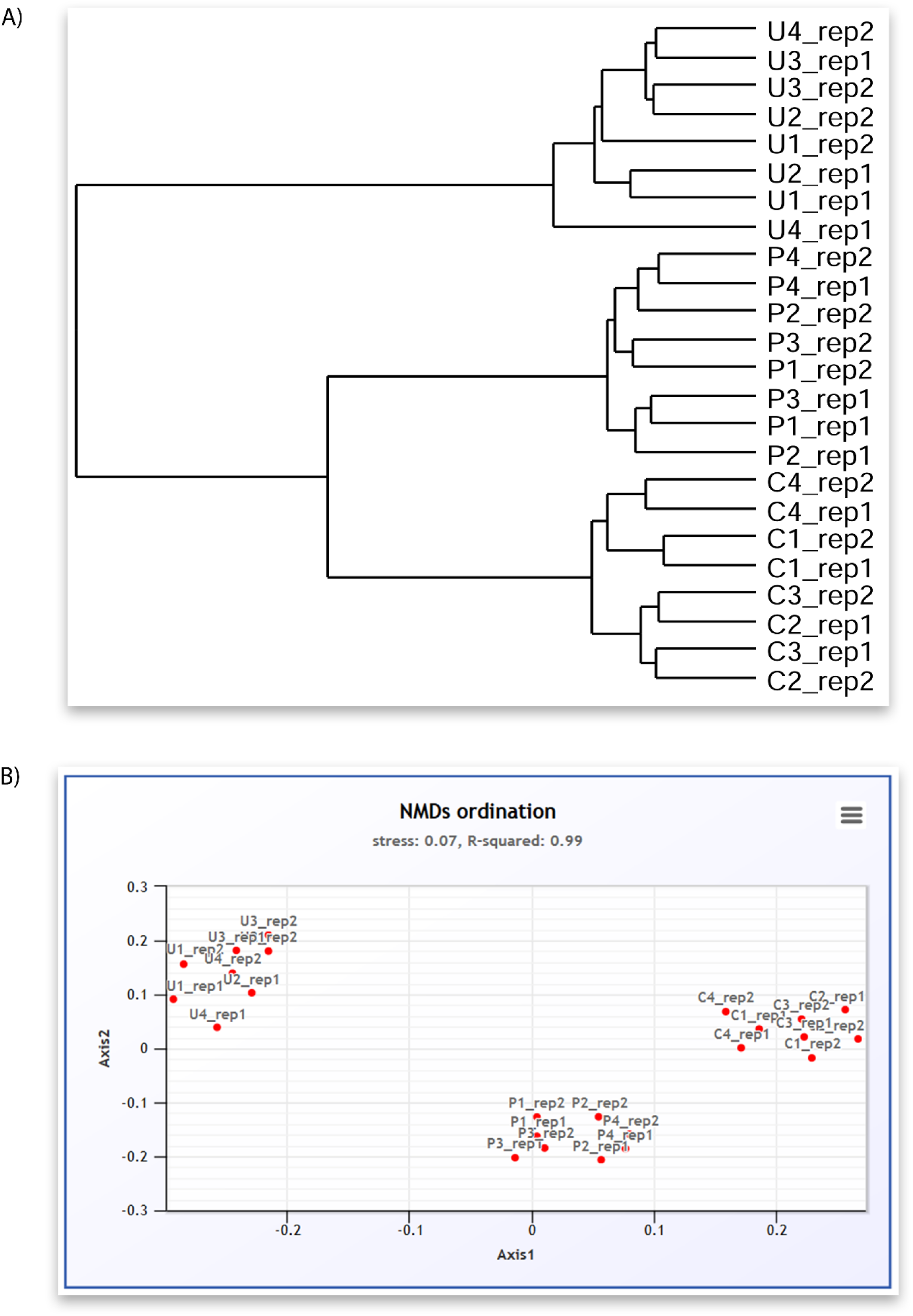
A screenshot montage of MetaAmp output in different view. A) Mock sample relation dendrogram describes the similarity of the mock samples to each other measured by Bray-Curtis dissimilarity distance. B) Mock sample NMDS analysis based on the Bray-Curtis dissimilarity distance.

Based on Bray-Curtis and ThetaYC dissimilarity matrixes, MetaAmp generated dendrograms to describe the similarity of the samples to each other. The trees showed that each type of the mock community clustered into a distinct branch (Figure 2A). Weighted UniFrac testing (P<0.01) confirmed that the three types of mock communities were significantly different. Principal coordinates and non-metric multidimensional scaling visualizations of the rarefied samples demonstrated a clear separation among the different types of mock communities (Figure 2B). The AMOVA testing results (P<0.01) further confirmed that the observed separation among the different types of communities was statistically-significant.

## Discussion

MetaAmp was shown to integrate available tools and reference databases for the comprehensive analysis of amplicon data. The easy-to-use yet versatile web interface allows users to access the pipeline from anywhere. At the same time, it also gives users the flexibility to configure the pipeline at different stages of the analysis. One limitation that MetaAmp shares with all other currently available tools is that the accuracy of the OTU taxonomic classification heavily depends on the underlying reference sequences/database. In the case of MetaAmp, taxonomic classification is based on the “Classify.seqs” command in Mothur, which can, as we have shown in this study, produce inaccurate taxonomic assignments when the targeted taxonomy is missing from the reference templates.

One challenge for all amplicon sequencing studies, which can be at least partially addressed with MetaAmp, is the problem of cross-talk within and between sequencing runs. This problem, resulting from sequencing and/or image analysis errors during the index sequencing phase of the MiSeq run (a separate step in the sequencing process), causes a small fraction of amplicons from one library to be incorrectly assigned to an index of another library (Nelson et. Al. 2014). In the mock community analysis we clearly showed the relevance of this problem by identification of 2-7 OTUs that were contaminants from DNA extraction kits and cross-talk. By analyzing all samples from a MiSeq run with MetaAmp, potential cross-talk can be identified. We strongly recommend the use of this procedure because our analysis showed that the contamination rate can be more than 2% in a MiSeq run. Within-run cross-talk can artificially inflate OTU numbers and diversity measurements if not properly addressed, leading to incorrect interpretation of results when investigating low abundance OTUs (Nelson et.al. 2014).

In conclusion, we presented and validated MetaAmp, a push-of-a-button pipeline for entry level users of amplicon sequencing. MetaAmp enables these users to rapidly generate solid results, addressing the basic questions of their studies. For advanced users, it offers convenient automation of the first steps of the analysis, preparing files that could be used as the starting point for more sophisticated tools. Since the launch of the website two years ago, it has already been used >2,800 times by scientists worldwide, which clearly indicates a strong demand for a simple tool of this type.

## Data Deposition

The mock community 16S rRNA gene amplicon sequencing data is available from the European Nucleotide Archive with run accession numbers ERR1938377 to ERR1938400 within study PRJEB19901 (http://www.ebi.ac.uk/ena/data/view/PRJEB19901). It also available to download from MetaAmp website.

## Author contributions

MetaAmp was conceived by XD and MS. MK contributed mock community creation and validation with input from CES, ET, DL and CL. CL contributed16S rRNA gene amplicon library preparation and sequencing on the MiSeq. The manuscript was written by XD and MS with input from all co-authors.

